# Degeneracy in Astrocytic Potassium Buffering: A Minimal Model Capturing the Interplay Between Local and Long-Range Mechanisms

**DOI:** 10.1101/2025.07.20.665757

**Authors:** Aitakin Ezzati, Nariman Kiani, Anton Ivanov, Jirsa Viktor, Damien Depannemaecker, Christophe Bernard

**Author notes:** co-last author (Damien Depannemaecker, Christophe Bernard).

## Abstract

Maintaining extracellular potassium (K+) homeostasis is critical for neuronal function, and astrocytes achieve this through a combination of local uptake and long-range spatial buffering. While degeneracy—the ability of different mechanisms to achieve the same function—is a fundamental property of biological systems, its role in astrocytic potassium buffering has remained unexplored.

We present a minimal mathematical model that identifies essential buffering mechanisms while ensuring tractability and interpretability. Incorporating Kir channels and gap junction coupling, the model reproduces experimentally observed astrocyte membrane dynamics under various pharmacological conditions

Parameter exploration reveals two levels of degeneracy. At the single-cell level, multiple parameter configurations yield similar membrane potential dynamics, indicating flexibility in local and spatial buffering contributions. At the functional level, despite variations in astrocyte morphology and buffering efficiency, homeostasis of extracellular K+ is restored, demonstrating homeostatic degeneracy.

These findings highlight the robustness of astrocytic potassium regulation, showing that diverse buffering strategies ensure stability. Our work establishes a theoretical framework for understanding how astrocytic heterogeneity contributes to robust ionic homeostasis and offers perspectives for studying pathological conditions where buffering mechanisms are impaired.

## 1 Introduction

Astrocytes, the most abundant glial cells in the central nervous system, are essential for maintaining neuronal homeostasis and supporting various neural functions [41]. These star-shaped cells are involved in many processes, including maintaining the blood-brain barrier, modulation of synaptic transmission through gliotransmission, and participating in neurovascular coupling, linking neuronal activity to blood flow. Astrocytes also play a critical role in neurogenesis and synapse formation, as well as in the CNS’s response to injury through reactive astrogliosis [36]. One of their most important functions is the regulation of extracellular ion concentrations, particularly potassium. The potassium ion (*K*^+^) plays a crucial role in the central nervous system, even small changes in its concentration can have a significant impact on neuronal excitability [5, 37]. Paradoxically, neuronal activity contributes to *K*^+^ accumulation in the extracellular space, necessitating extra-neuronal mechanisms to maintain *K*^+^ homeostasis. Failure of these clearance mechanisms has been implicated in pathologies such as epilepsy [12]. By efficiently buffering extracellular potassium ions, astrocytes prevent the hyperexcitability of neurons. Multiple mechanisms participate to this buffering process, including direct uptake via the astrocytic Na+/K+-ATPase (NKA), which is particularly sensitive to extracellular *K*^+^ concentrations [21].

Spatial buffering involves astrocytes using their extensive interconnected network, linked by gap junctions composed of connexin43 and connexin30, along with inward-rectifying potassium channels (Kir4.1), to redistribute *K*^+^ ions. This process transfers excess potassium from regions of high neuronal activity to areas where levels are more balanced, helping maintain ionic homeostasis [26, 30]. The membrane of mature protoplasmic astrocytes is highly permeable to *K*^+^, characterized by numerous channels that constitute the majority of astrocytic conductances [47]. This permeability, combined with extensive gap junction coupling, imparts isopotentiality to the astrocytic network, where neighboring coupled astrocytes contribute to the passive current profile via their junctional conductance [24]. Despite extensive research, our understanding of the interactions between individual astrocytes and extracellular ionic compositions remains incomplete. This knowledge gap is primarily attributed to the limited biophysical data available for astrocytes, a consequence of their electrical passivity and the technical difficulties associated with isolating specific ionic currents using conventional electrophysiological methods. Furthermore, the dynamical nature of these processes adds another layer of complexity to studying astrocytic functions in ionic regulation.

In the context of epilepsy, core astrocytic mechanisms such as Kir4.1 channels and connexin-mediated coupling have been particularly well-characterized, highlighting their critical roles in potassium buffering [13, 14]. Nevertheless, several important questions remain unanswered. The dynamic regulation of Kir4.1 expression during different epileptic phases and the optimal balance of gap junction coupling needed to maintain effective potassium clearance without provoking excessive neuronal synchronization continue to be debated [6, 25, 29]. Additionally, significant uncertainties persist regarding regional differences in buffering efficiency, notably between hippocampal and cortical astrocytes [9, 38, 44], as well as the impacts of neuroinflammatory and neuromodulatory factors on potassium homeostasis [7]. Addressing these gaps necessitates advanced computational models that accurately capture the intricate dynamics of astrocytic potassium regulation, providing deeper insights into their physiological and pathological roles and informing potential therapeutic strategies [11, 28]. Given the complexity of astrocytic functions, spanning metabolic, homeostatic, and neuromodulatory roles, and their highly plastic morphology and spatiotemporal variability, modeling becomes an invaluable tool for testing hypotheses and understanding system dynamics. The reductionist approach in modeling, emphasizing the capture of essential dynamics with a minimal number of variables, has proven effective in neuroscience by offering a simplified framework for understanding the interplay of mechanisms. [33, 34].

In this study, we aim to create a simple reduced model for astrocyte membrane dynamics, with a focus on both short- and long-range potassium buffering capabilities. Our approach is based on the idea that, despite the variability in astrocytic properties, certain dynamical features of cell membranes remain consistent. We expect that different mechanisms can produce the same observed dynamics, a concept known as degeneracy. To explore this, we developed a model that incorporates key elements of astrocytes involved in [*K*^+^] dynamics, as identified in the literature, and validated it against experimental data. Using parameter optimization techniques, we were able to identify a range of parameter sets that produce similar observed dynamics, confirming degeneracy. This study not only sheds light on the mechanisms of astrocytic potassium buffering but also highlights the effectiveness of minimal modeling in capturing complex biological processes. By uncovering potential degeneracy in astrocytic functions, our work enhances our understanding of the robustness and adaptability of these critical central nervous system components.

## 2 Materials and Methods

This project aimed to develop a minimal biophysical model of astrocyte function to investigate its role in maintaining ionic homeostasis within the interstitial space. The model was designed to incorporate only the essential mechanisms contributing to potassium (*K*^+^) buffering capacity, as identified through experimental electrophysiological studies. By focusing on a reduced set of fundamental processes, we sought to capture the core dynamics underlying astrocytic *K*^+^ regulation, providing a simplified but dynamically accurate representation of their contribution to ionic homeostasis in the central nervous system.

### 2.1 Ex vivo experiments: Brain slice preparation

Male FVB mice, anesthetized with sevoflurane, were decapitated. The brain was swiftly removed from the skull and submerged in ice-cold artificial cerebrospinal fluid (ACSF) (in mM): NaCl 126mM, KCl 3.50 mM, NaH_2_PO_4_ 1.25 mM, NaHCO_3_ 25mM, CaCl_2_ 2mM, MgCl_2_ 1.3mM, and glucose 10mM, with a pH value maintained between 7.3 and 7.4. The ACSF was aerated with a 95% O_2_/5% CO_2_ gas mixture. Using a tissue slicer (Leica VT 1200s, Leica Microsystem, Germany), coronal slices measuring 300 m were cut. Before slicing, the brain was immersed in an ice-cold cutting solution (in mM): K-gluconate 140mM, HEPES 10mM, Na-gluconate 15mM, EGTA 0.2mM, and NaCl 4mM, with the pH adjusted to 7.2 with KOH. Subsequently, to preserve the surface from damage resultant from slicing, the slices were incubated for 15 to 20 minutes at room temperature in a choline chloride solution (in mM): Choline Chloride 110mM, KCl 2.5mM, NaH_2_PO_4_ 1.25mM, MgCl_2_ 10mM, CaCl_2_ 0.5mM, NaHCO_3_ 25mM, Glucose 10mM, Na-Pyruvate 5mM, constantly bubbled with a 95% O_2_/5% CO_2_ gas mixture. To label the astrocytes, the slices were incubated for 20 minutes in a modified ACSF solution containing 1M of sulforhodamine 101 (SR101, Merk), a fluorescent dye specifically absorbed by astrocytes and not neurons. The slices were allowed to recover in a dual-side perfusion holding chamber with continuously circulating ACSF for at least one hour before experimentation.

### 2.2 Electrophysiological recording of the astrocytes

For the electrophysiological recordings, slices were immersed in a low-volume (2 mL) recording chamber and continuously perfused with ACSF at 32°C and a 5 mL/min perfusion rate. Astrocytes in the stratum radiatum of the CA1 region of the hippocampus were identified using excitation/emission light wavelengths of 586/605 nm. Patch pipettes were fabricated from borosilicate glass tubing (1.5 mm outer diameter, 0.5 mm wall thickness) and filled with an intrapipette solution (in mM): 20 KCl, 115 K-gluconate, 10 HEPES, 1.1 EGTA, 4 MgATP, 10 Na-phosphocreatine, and 0.4 Na2GTP. Signals were transmitted to a Multiclamp 700A amplifier (Molecular Devices), digitized at a rate of 10 kHz using a DigiData 1550 interface (Molecular Devices) connected to a personal computer, and analyzed using ClampFit software (Molecular Devices). The membrane potential of astrocytes was measured using the patch-clamp technique, whole-cell configuration, in the current clamp mode. *K*^+^ delivery. The iontronic micropipette was connected to the second head-stage. A chlorinated silver wire facilitated the transfer of current from the head stage to the reservoir, filled with 100 mM KCl. The protocol has already been established in previous work [3]. In brief, the iontronic pump operated in current clamp mode, allowing driving current control. *K*^+^ delivery to the patched astrocyte. *K*^+^ delivery was performed in 6 trials, beginning with a 50 nA current drive and increasing by 25 nA per trial. Each trial consisted of a 20-second baseline, a 20-second drive, and a 30-second recovery period. In experiments involving blockers, a round of 6 trials was designated for ACSF containing meclofenamic acid, MFA (0.1 mM) a blocker of gap junctions in the astrocytes, followed by another round of trials with MFA-containing ACSF supplemented with 0.1 mM BaCl2, a blocker of inward-rectifying *K*^+^ channels (Kir4.1).

### 2.3 Calibration of the iontronic [*K*^+^] delivery pump

Potassium-selective microelectrodes were fabricated following the procedure outlined by Lux and Neher (1973) [23]. In summary, electrodes were pulled from double-barrel theta glass (TG150-4, Warner Instruments, Hamden, CT, USA). The reference barrel was filled with a 154 mM NaCl solution. The silanized ion-sensitive barrel tip (prepared with 5% trimethyl-1-chlorosilane in dichloromethane) was filled with a potassium ionophore I cocktail A (60031 Fluka, distributed by Sigma-Aldrich, Lyon, France) and backfilled with 100 mM KCl. Measurements of K^+^-dependent potentials were conducted using a high-impedance differential DC amplifier equipped with negative capacitance feedback control. This setup enabled compensation for the microelectrode capacitance necessary for recording rapid changes in extracellular K^+^ [K^+^]_ext_. The electrodes were calibrated before each experiment. Pulses lasting 7 seconds with a 30-second interval between pulses (from the onset of one to the onset of the next) were utilized. The iontronic pump was placed at the immediate vicinity of the K^+^ sensor, less than 5 *µ*m, inside the recording chamber containing only ACSF and was driven in 5 nA increments, commencing from 5 nA and culminating at 190 nA. The steady-state value from each trial was fitted in the calibration monoexponential fitting to obtain the resulting K^+^ concentration, which was then plotted against the drive. The relationship between the K^+^ from the iontronic pump and the pump current drive had a linear fit (R^2^ = 0.99).

### 2.4 Syncytial Astrocyte Model

We considered a model of a single astrocyte connected via gap junction to other astrocytes through the syncytium that is assumed to be in the steady state, as done in previous work [17].

As shown in the schematic in Fig. 1, on one hand, the astrocyte is surrounded by the extracellular space (ECS) and connected to the syncytium through gap junctions. The ECS, on the other hand, is coupled to an infinite *K*^+^ reservoir which resembles the long-range diffusion in the tissue or the bath in slice experiments. In this configuration, the ECS and syncytium are only connected indirectly via astrocytic transmembrane currents. This indirect connection is reflected in the mass conservation equation(4).

**Figure 1:**
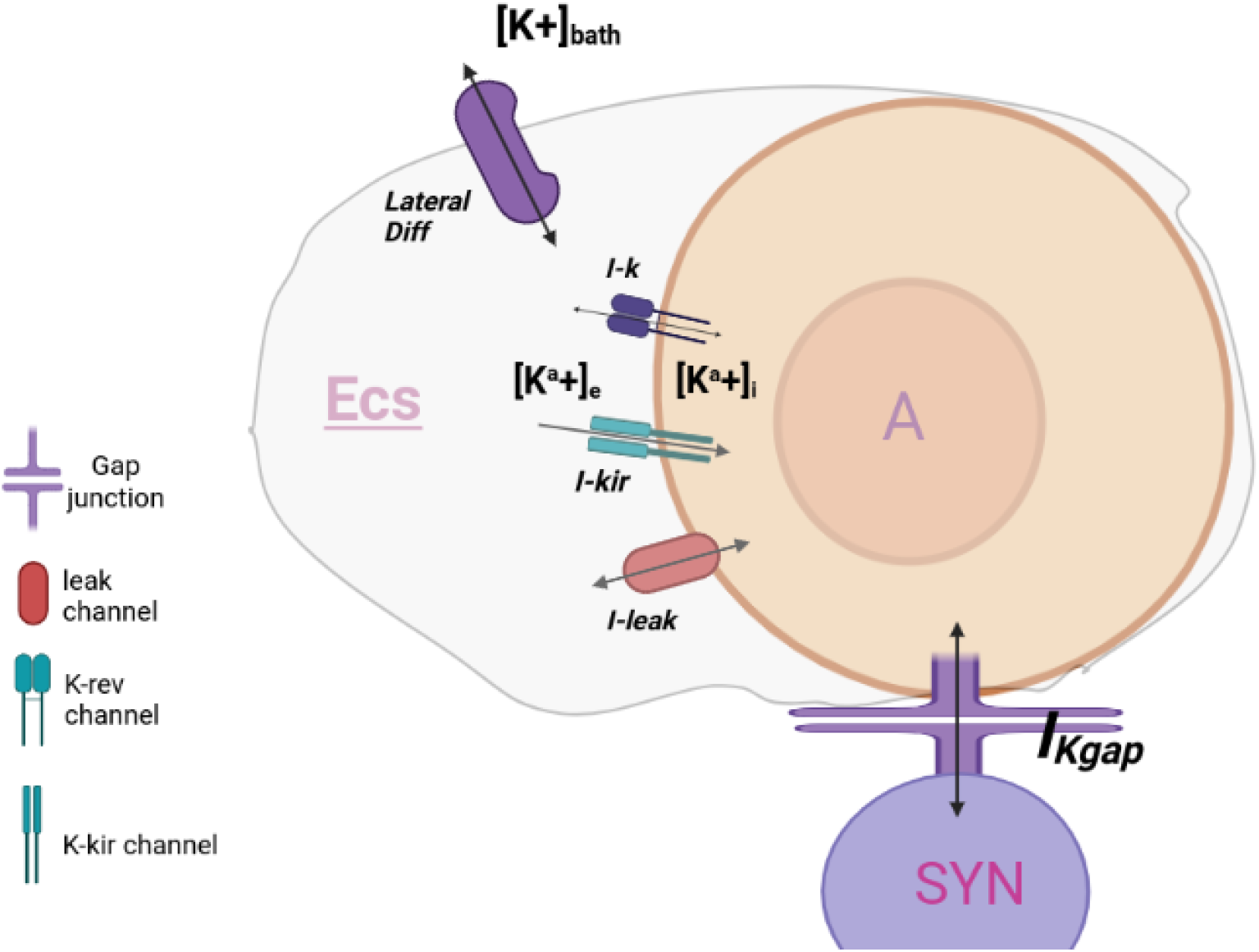
A single astrocyte compartment (A) is connected to the syncytium (Syn) via a gap junction (Gap) and interfaces with the extracellular space (ECS) through three types of transmembrane conductances: a general leak, a passive K^+^ channel (K_rev_), and the Kir4.1 channel. The ECS is coupled to a potassium reservoir (Bath) to account for lateral diffusion. This configuration ensures ionic mass conservation across compartments while enabling the study of short- and long-range K^+^ buffering dynamics.

Incorporating *K*^+^ channels and the gap junction connection replicates biophysical features such as high *K*^+^ permeability and hyperpolarized membrane. These are the main mechanisms behind passive *K*^+^ conductance. According to [47], the main functional implication of this passivity is a strong buffering capacity. Although this model does not consider the three-dimensional space, it instead relies on the biophysical properties mentioned above as phenomena, we could effectively incorporate different modalities of *K*^+^ buffering in this model, namely local uptake, lateral diffusion in ECS, and intracellular electrical coupling, thus spatial buffering. The dynamics of the astrocyte membrane potential is modeled by the following differential equation:

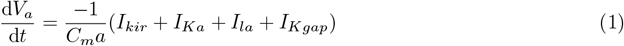

Due to the high permeability of astrocytic membrane to *K*^+^ compared to other ions, we decided to only consider the effect of *K*^+^ dynamics on the variation of the membrane potential as it has been done before, [19], by including the following currents:

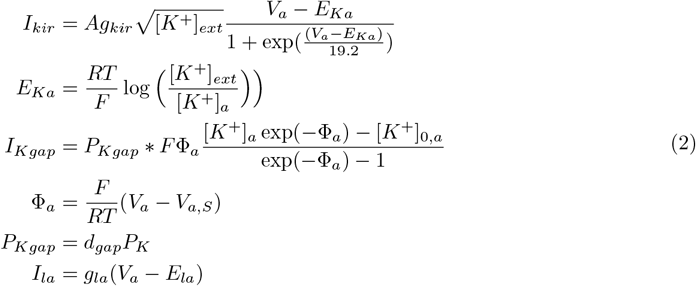

The *I*_*kir*_ is the inward rectifying current that is modeled as in Ransom et al. [31]. The A parameter is a scaling factor of dimension 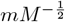. *E*_*Ka*_ is the reversal potential of *K*^+^. To have access to some biophysical parameters such as coupling strength, we resorted to a more precise description using Goldman-Hodgkin-Katz formalism for gap junction current *I*_*Kgap*_, as it was done previously in [24]. This formulation includes the explicit term for averaged permeability over the cell *P*_*Kgap*_ = *d*_*gap*_*P*_*K*_. As it was shown in [17,40] the overall permeability depends on the number of connected astrocytes and the coupling coefficient. Here *d*_*gap*_ is de fined as the coupling strength, which can be a proxy for these values. (check this further). This nonlinear function increases as a function of both the potential difference and the concentration gradient between the astrocyte and the syncytium. It integrates elements of ionic coupling and isopotentiality, contributing to the spatial buffering of ions. *I*_*la*_ is a general leak current with a small conductance to compensate for the contribution of *K*^+^ and *Na*^+^ leaks that are not explicitly modeled as a common practice in other models previously described. [32].

#### Ionic concentrationn

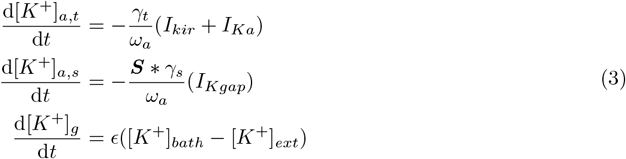

The intracellular concentration change, Δ[*K*^+^]_*a*_ is due to the contribution of the transmembrane mechanisms at the side of the ECS and gap junctional coupling. Here, we decided to separate each contribution as an independent variable as Δ[*K*^+^]_*a,t*_ and [*K*^+^]_*a,s*_ respectively, as each of them is in direct contact with a different compartment. To convert the rate of concentration change to current, as explained in Cressman et al. [10], we use a conversion factor 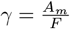 divided by the volume of the whole cell *ω*_*a*_. Note that since we are using *m* as the morphological unit while Molar for concentrations, we need to multiply the right-hand side of these equations by 10 to correct for the unit incompatibility [15].

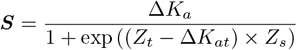

The function ***S*** represents a sigmoïd function that models the threshold-dependent behavior of the astrocytic spatial buffering of *K*^+^. This formulation is designed to capture the adaptability of spatial buffering engagement in response to varying extracellular *K*^+^ concentrations. The incorporation of this function was motivated by recent findings from Bellot-Saez et al. [7], which demonstrated that the rate of extracellular *K*^+^ clearance by astrocytes exhibits a non-linear dependence on *K*^+^concentration. Specifically, their observations revealed that *K*^+^ clearance becomes more rapid and efficient at higher extracellular *K*^+^ concentrations, suggesting an adaptive mechanism for *K*^+^ homeostasis.

According to their experiments involving blocking Kir4.1 channel and gap junctions, they concluded that for the low concentrations, only the local uptake mechanisms are effectively involved, and after the accumulation of high concentrations, such as above 10 mM, the spatial buffering is engaged effectively, which makes the clearance rate much faster. Their observations also indicated that blocking Kir 4.1 effectively hampers the efficacy of the spatial buffering. Previous literature shows that blocking Kir channel currents by Ba+ is effectively possible when gap junctions are blocked as well [35], [44].

Hence, to capture the interdependence between these two mechanisms, we needed to introduce a function that can limit the capacity of gap junctions, which hampers but does not abolish the local uptake mechanism. Furthermore, the function should detect the accumulation of *K*^+^ as a threshold for switching to the full capacity of the gap junction. Therefore, a sigmoid that could introduce flexible thresholds depending on its parameters is the most suitable choice. Here, *Z*_*t*_ represents the threshold for intracellular flux to accumulate and *Z*_*s*_ controls the degree to which the spatial buffering is limited before the threshold is reached. These values could depend on many factors such as experimental conditions, specific characteristics of the astrocyte phenotype, which varies depending on the region in the brain, as well as temporal variability due to circadian and multidian cycles [27, 42]. Additionally, according to Buskila et al. [7] the presence of some neuromodulators, such as norepinephrine and Dopamine, to name a few, has been shown to control the level of gap junction coupling and thus, via modulating the speed of buffering, effectively shape the dynamics of extracellular *K*^+^. Therefore, it is safe to say that our formulation captures a certain bidirectional aspect of neuron-astrocyte interaction.

Since the gap junction is only indirectly connected to the ECS (as shown in the schematic 1) via trans-membrane currents, the logical choice was to set a threshold based on Δ[*K*^+^]_*t*_ and not [*K*^+^]_*o*_. Therefore, the accumulation of *K*^+^ inside the astrocyte is a proxy for accumulation of *K*^+^ in the ECS in this formulation.

To clarify this further, we have to describe the forth equation, which represents the diffusion of *K*^+^ in or out of the ECS to-from the bath depending on the difference between the two concentrations, and, diffusion coefficient*ϵ*. Ergo, the competition between the influx from the bath *K*_*g*_ and the change of astrocyte interacellular potassium, Δ[*K*^+^]_*a*_, rise up to the change in Δ[*K*^+^]_*o*_.

The magnitude of the rate of change in spatial siphoning [*K*^+^]_*s*_, intuitively relates to the change in intracellular concentration Δ[*K*^+^]_*a*_. This guaranties that the steady state is at the point where Δ[*K*^+^]_*t*_ = −[*K*^+^]_*s*_,. Its worth mentioning that here we are dealing with an open system which can have a constant flow between sinks and sources as the condition for steady states. [*K*^+^]_*t*_, [*K*^+^]_*s*_ and [*K*^+^]_*g*_ all are associated with the change of concentration. Therefore, as long as the rate of change of these variables amount to a constant value, be it 0 or not, that could satisfy the condition of zero sum total currents in and out of membrane. The following mass conservation equations match the model configuration.

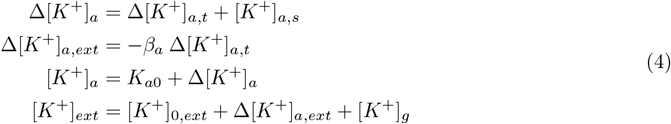

These relationships define the constraints on *K*^+^ movement between compartments, ensuring proper accounting of their inflow and outflow within the system.

Since we only considered the change in the concentration of *K*^+^, a cation, in order to ensure electroneutrality in this system, we added a general leak term which can be considered as the average leak over the whole membrane for other ion species such as *Na*^+^ and *Cl*^*−*^. These currents could provide the counterbalance needed to guarantee electroneutrality.

##### 2.4.1 Optimization algorithm

To find the sets of parameters leading to the observed dynamics in each experiment, we utilized an automatic hyperparameter optimization framework, Optuna [1]. It primarily uses the Tree-structured Parzen Estimator (TPE) approach, which models the search space probabilistically using Bayesian optimization principles. It is particularly effective because it can quickly converge to the optimal parameters by prioritizing hyperparameter configurations that appear more promising.

For calculating the time constant, *τ*_*rise−decay*_ we employed curve-fit as in [39] well as optuna to find the best exponential fit for the 10-90(90-10) percent [7] of the rise and decay section of the traces. The decay time constant (*τ* ) was determined by fitting the membrane potential traces to a single-exponential decay function of the form:

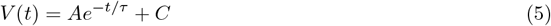

where *A* is the amplitude, *τ* is the decay time constant, and *C* represents the steady-state offset. The fitting was performed using nonlinear least-squares optimization (`curve_fit` from SciPy) on the post-stimulation segment of the traces, when the iontronic pump stops.

**Table 1:** Model parameters and their definitions. List of an example parameter configuration within biophysical range, used in the syncytial astrocyte model, including membrane conductances, ionic concentrations, coupling strengths, and bath properties. Units and brief descriptions are provided to clarify each parameter’s role in governing transmembrane fluxes and compartmental interactions.

| Name | Value and Unit | Description |
| --- | --- | --- |
| **Astrocyte:** |  |  |
| $C_a$ | $100 \mu\text{F}/\text{cm}^2$ | Membrane capacitance |
| $g_{Kir}$ | $10 \text{ mS}/\text{cm}^2$ | Maximal potassium conductance of Kir channel |
| $A$ | $1 \text{ mM}^{-\frac{1}{2}}$ | Pseudo parameter |
| $g_{la}$ | $0.37 \text{ mS}/\text{cm}^2$ | General leak conductance |
| $\omega_a$ | $2160 \mu\text{m}^3$ | Extracellular volume |
| $\gamma_t$ | $5 \mu\text{m}^2 \text{mol}^{-1} \text{C}^{-1}$ | Conversion factor |
| $\gamma_s$ | $6 \mu\text{m}^2 \text{mol}^{-1} \text{C}^{-1}$ | Conversion factor |
| $\omega_a$ | $2000 \mu\text{m}^3$ | Volume |
| $\omega_o$ | $720 \mu\text{m}^3$ | Volume |
| $\beta_a = \omega_a/\omega_{ext}$ | 2.77 | Intra/extracellular volume ratio |
| $\sigma_a$ | $1600 \mu\text{m}^2$ | Surface |
| $P_k$ | $4.8 \times 10^{-5} \text{ cm}/\text{s}$ | Membrane potassium permeability |
| $d$ | 1 | Gap junction coupling strength |
| $F$ | $96458.0 \text{ C}/\text{mol}$ | Faraday constant |
| $RT$ | $0.593 \text{ Kcal}/\text{mol}$ | $Gasconstant \times Absolutetemperature$ |
| $\epsilon$ | $10^{-3} \text{ m}^{-1} \text{s}^{-1}$ | Diffusion coefficient in bath |
| $V_{a,l}$ | -70 mV | Reversal leak potential |
| $V_s$ | -90 mV | Steady-state syncytium membrane potential |

## 3 Results

The results are organized in two main parts: (1) the underlying dynamics and validation against experimental data, and (2) the exploration of parameter space to characterize degeneracy in astrocytic potassium buffering.

### 3.1 Kinetics of K+ buffering

Our model captures the response of astrocytes to changes in extracellular potassium concentration through the parameter *ϵ* (s^*−*1^), which controls the diffusion of potassium from an infinite source bath (equation (3)). The initial condition of the simulation for the unperturbed system is at *K*_Bath_ = *K*_*o*_. To probe the effect of elevated levels of K^+^ on the dynamics, we set *K*_Bath_ to a fixed higher value for the duration of the simulation.

Figure 2 illustrates the temporal evolution of membrane potential (*V*_*a*_), transmembrane K^+^ flux (*K*_*t*_), and intracellular K^+^ concentration (*K*_*a*_) in response to an increase in *K*_Bath_ as a function of different *ϵ* values. The time series reveals a clear separation between fast and slow transients. The membrane potential (*V*_*a*_) changes rapidly, while the transmembrane potassium flux (*K*_*a,t*_) evolves more slowly. The intracellular potassium concentration (*K*_*a*_) contains both a fast and a slow component. Increasing *ϵ* accelerates both the fast and slow kinetics.

**Figure 2:**
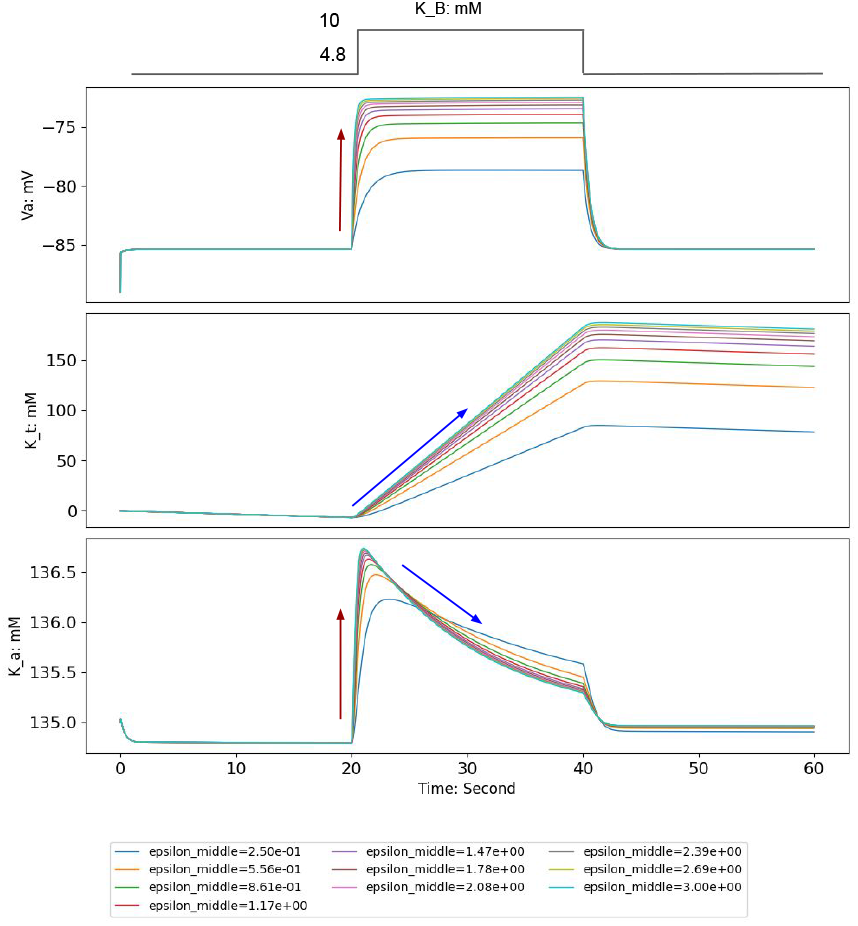
Astrocyte dynamics under varying levels of local *K*^+^ input reveal distinct temporal profiles. Simulation results from a fixed parameter configuration illustrate how changes in extracellular potassium concentration—driven by varying the intensity of *K*^+^ input from the bath (parameter *ϵ*)—affect astrocyte dynamics. Each panel shows the temporal evolution of a key variable: membrane potential (*V*_*a*_, top), transmembrane potassium flux (*K*_*t*_, middle), and intracellular potassium concentration (*K*_*a*_, bottom). Colored traces correspond to increasing values of *ϵ*, as indicated in the legend. Higher input levels result in faster dynamics across all variables. The membrane potential (*V*_*a*_) depolarizes rapidly; *K*_*a*_ displays a biphasic profile with a fast rise followed by a slower decay; and *K*_*t*_ evolves more gradually. Red arrows highlight the fast rise in *V*_*a*_ and *K*_*a*_, while blue arrows indicate the slower dynamics of *K*_*t*_ and the slow decay of *K*_*a*_.

The rate of change in intracellular potassium concentration (*K*_*a*_) is governed by equations involving the local potassium current through Kir channels (*I*_Kir_) and the gap junctional potassium current (*I*_*k*,gap_). The parameters controlling these currents are the maximum potassium conductance of Kir channels (*g*_Kir_), the membrane potassium permeability (*p*_*k*_), and the gap junction coupling strength (*d*).

For instance, increasing *g*_Kir_ enhances the astrocyte’s ability to rapidly uptake extracellular potassium, thereby accelerating the initial response to potassium surges. Similarly, variations in *p*_*k*_ affect the membrane’s permeability, modulating the rate at which potassium ions cross the membrane, while adjustments in *d* influence the extent of potassium redistribution through gap junctions, affecting spatial buffering capacity. Thus, these parameters directly control the speed and efficiency of potassium clear-ance, shaping the balance between fast transients and slower, sustained adaptations in astrocyte function. This interplay of parameters reflects the system’s capacity to adapt to varying physiological conditions, highlighting the critical role of biophysical processes in modulating the astrocyte’s temporal dynamics.

This timescale separation allows the system to flexibly adjust its response to external stimuli by balancing immediate transients with longer-term adaptations. For instance, rapid changes in membrane potential can quickly modulate ionic currents, while slower adjustments in potassium fluxes through channels and gap junctions sustain prolonged responses [18].

According to the literature [7, 44], astrocytes exhibit both slow and fast potassium buffering modes that become dominant at lower or higher extracellular K^+^ concentrations, respectively. The slow mode is frequently attributed to local mechanisms such as Na^+^/K^+^-ATPase activity, whereas the fast mode primarily involves Kir channels in conjunction with gap junctions. Our model replicates these two modalities by including a sigmoidal gating function that modulates spatial buffering. This approach captures the essential effects of local buffering without explicitly modeling the Na^+^/K^+^-ATPase or additional exchangers, thereby keeping the model minimal and avoiding the need to track Na^+^ dynamics.

### 3.2 Experimental validation of the model

The validation of the model was treated as an optimization problem. We used the membrane potential recordings from patch-clamp experiments (see 2.2) as our target trajectories. The estimated range of extracellular K^+^ concentrations, derived from iontronic pump calibration (see 2.3), was used to define the interval for the *K*_Bath_ parameter in the optimization algorithm (see 2.4.1).

To validate our model, we compared simulated membrane potential traces with experimental recordings obtained under three conditions: control, meclofenamic acid (MFA, a gap junction blocker), and MFA plus barium chloride (which additionally blocks Kir4.1 channels).

To simulate the iontronic K^+^-pump experimental conditions (Fig 3a), we initialized the model with identical K^+^ concentrations in the bath and extracellular space (ECS). We then introduced the *K*_Bath_ hyperparameter elevation only during active iontronic pump delivery, so that diffusion from the bath would approximate the K^+^ delivery from the iontronic pump (Figure 3a).

**Figure 3:**
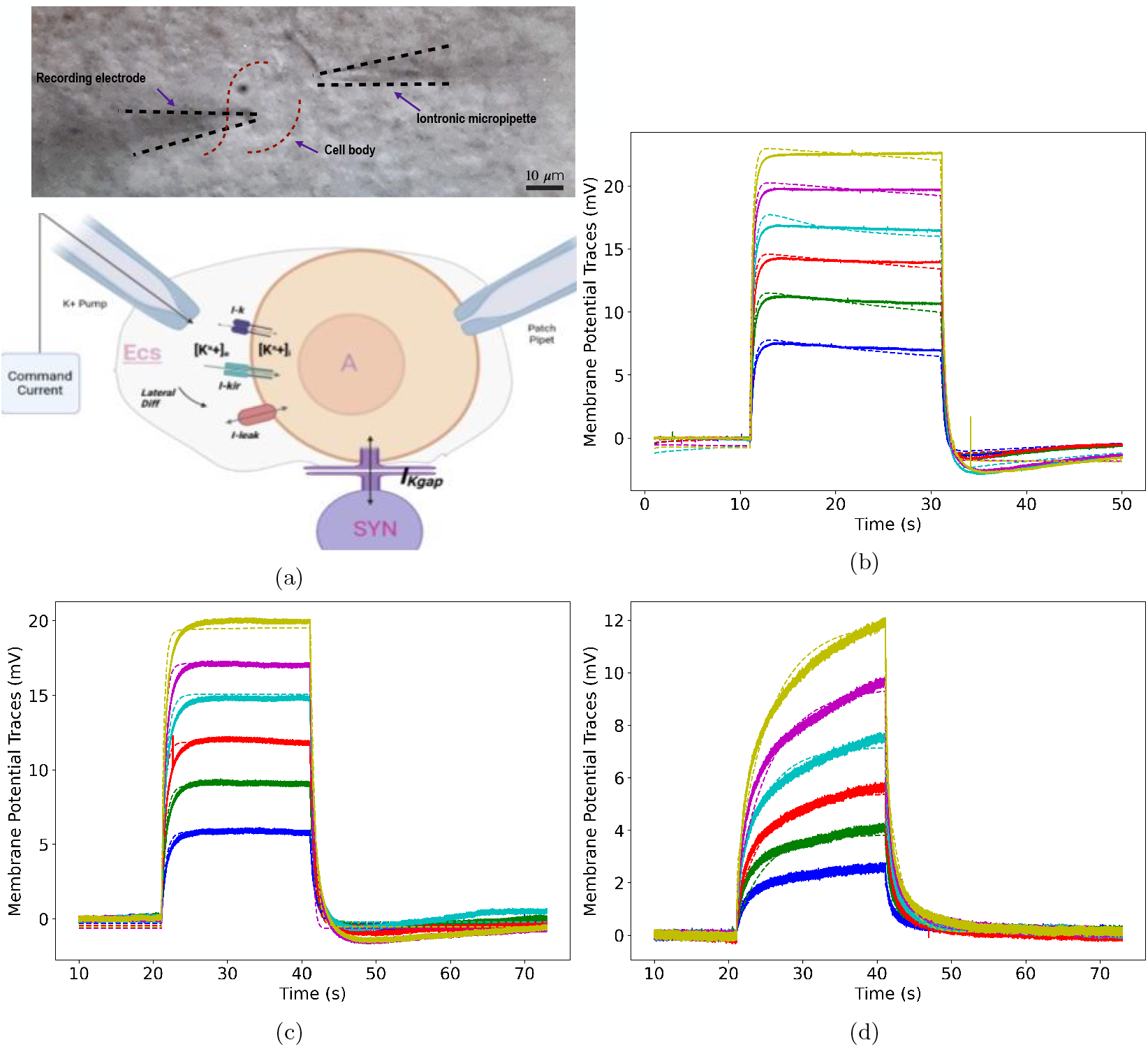
Reproducing experimental observations. (a) On the upper left, an image of the experimental setup featuring a patched astrocyte with a *K*^+^ iontronic pump positioned at a 10 *µM* distance. The bottom panel presents a schematic of the astrocyte model, illustrating the mechanisms incorporated under the iontronic pump experiment, noting that the *K*^+^ bath is replaced by the iontronic pump. Panels (b)control condition, (c) MFA condition: treatment with meclofenamic acid and (d) BA+ MFA condition: treatment with both meclofenamic acid and Barium, display membrane potential time series for each experimental condition. Each colored trace corresponds to a different iontronic pump current value (blue: 50, green: 75, red: 100, cyan: 125, purple: 150, yellow: 175). Solid lines represent the real, experimentally measured traces, compared with their simulated counterparts generated by the model.

We used six membrane potential traces recorded under each of the three experimental conditions—one for each increment of K^+^ delivery by the iontronic pump—as optimization targets. The parameter configurations resulting in the best fits for the experimental potential traces were achieved after 2500 converged trials of parameter search.

The model successfully reproduced key aspects of astrocyte membrane dynamics across all experimental conditions (Figure 3). Under control conditions (Figure 3b), we observed a slight downward slope during active iontronic pump stimulation that disappeared with MFA application. Both control and MFA conditions exhibited undershoots whose amplitudes increased with higher iontronic pump currents, with deeper troughs in control conditions suggesting enhanced buffering capacity due to gap junctions.

After adding barium to MFA treatment (Figure 3d), the undershoot disappeared for all traces, consistent with the impairment of gap junction coupling and blockage of Kir channels. Despite not including other buffering mechanisms such as the Na^+^/K^+^-ATPase pump, our model successfully reproduced these experimental results, suggesting that the basic mechanisms in our framework are sufficient to capture the key dynamics.

#### 3.2.1 Astrocyte membrane potential as a function of extracellular *K*^+^

The model correctly simulated the depolarizing effect of increased extracellular potassium on astrocyte membrane potential (Figure 4). We quantified this by calculating the amplitude of membrane potential increase across increments of iontronic pump currents for each experimental condition.

**Figure 4:**
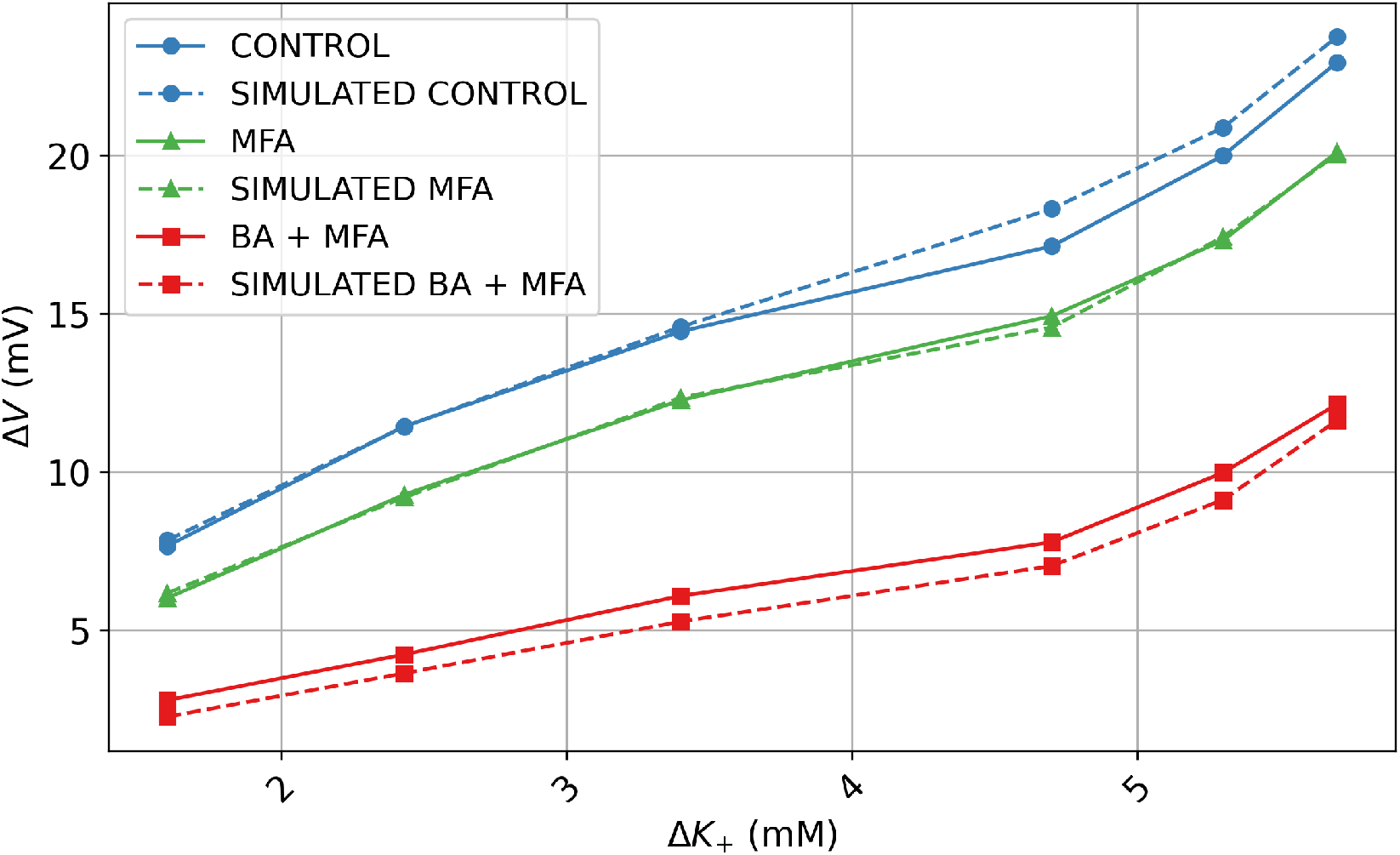
Amplitude of depolarization as a function of estimated potassium input. Experimental and simulated membrane potential amplitudes are plotted against Δ*K*^+^, the estimated increase in extracellular potassium concentration delivered by the iontronic pump. Data are shown for three pharmacological conditions: control, MFA, and BA+MFA. Solid lines represent experimental recordings, and dashed lines show the corresponding model simulations. Each point indicates the peak depolarization during the stimulation window for a given level of Δ*K*^+^.

After barium application, the lower yet increasing depolarization amplitude (Figure 4, red curve) can be attributed to the decrease in K^+^ gradient due to accumulation in the extracellular space. Under control conditions and MFA, the high K^+^ conductance led to greater depolarization amplitude, indicating stronger depolarizing currents compared to the barium condition.

The flux of K^+^ through the membrane is determined by the Nernst potential (*E*_*K*_), which depends upon the intracellular and extracellular ratio of K^+^. Increasing *K*_*a*_ (for a fixed *K*_ext_ = 5 mM) hyperpolarizes *E*_*K*_, while increasing *K*_ext_ (for a fixed *K*_*a*_ = 140 mM) depolarizes *E*_*K*_ (Figure 5, blue and green lines, respectively). Changing both concentrations while keeping a constant ratio does not change *E*_*K*_ (Figure 5, red line). The more biophysically probable scenarios are shown in dashed-dotted lines, where the *K*_*a*_ rate of change is either faster (red) or slower (orange) than that of *K*_ext_. The ratio was varied from 0.035 (solid red line) to the extreme cases (0.017 and 0.07, blue and green, respectively), including intermediate values (0.028–0.032, 0.038–0.043), shown as red and orange dotted and dashed lines.

**Figure 5:**
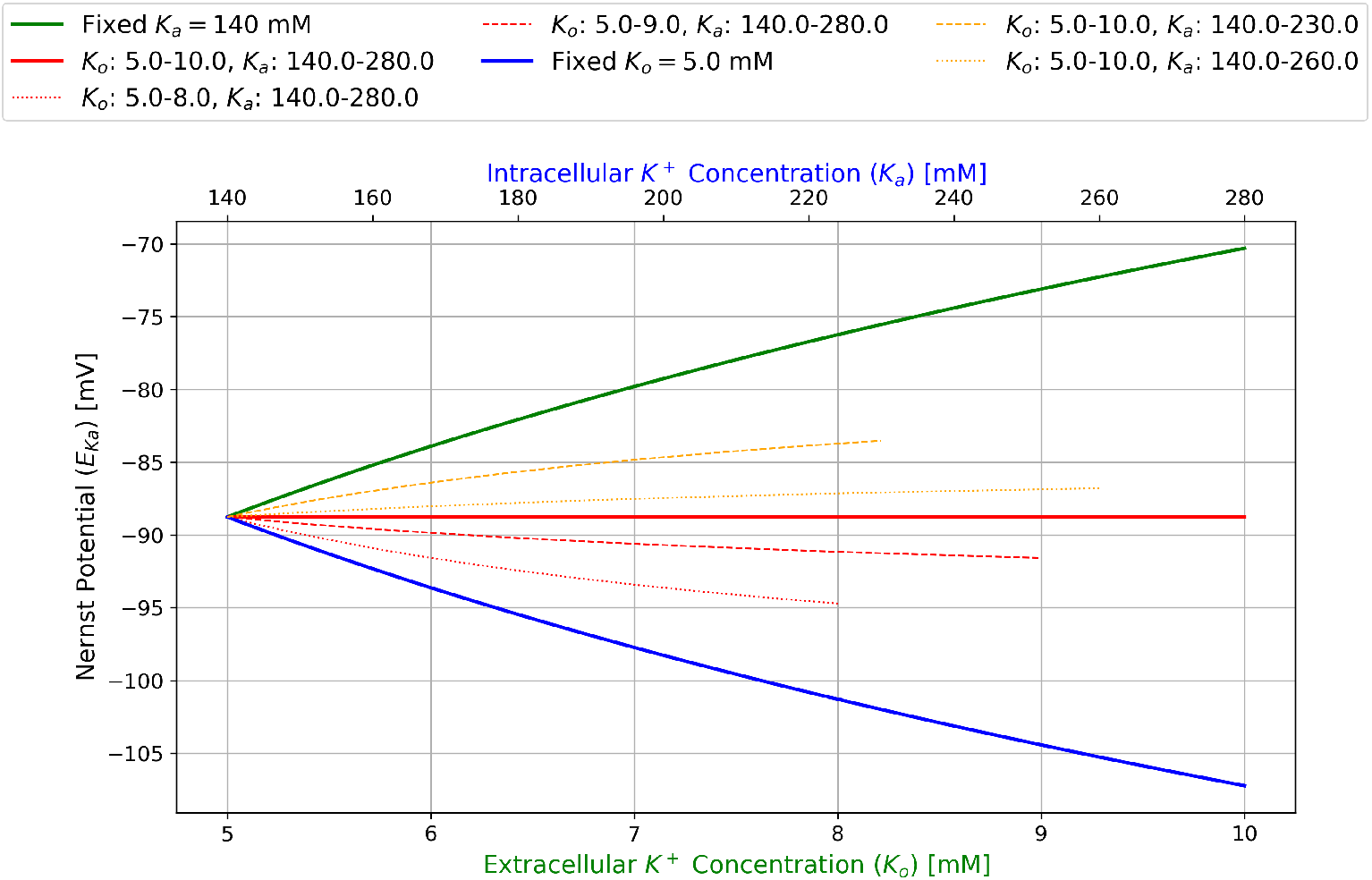
Reference map of reversal potential shifts induced by differential intra- and extra-cellular potassium dynamics. Each line shows the potassium reversal potential *E*_*K*_, computed using the Nernst equation, for different combinations of intracellular (*K*_*a*_) and extracellular (*K*_*o*_) potassium concentrations. The blue line shows *E*_*K*_ as *K*_*a*_ increases while *K*_*o*_ is held constant; the green line shows *E*_*K*_ as *K*_*o*_ increases while *K*_*a*_ is fixed. The red line corresponds to constant *K*_*o*_*/K*_*a*_, resulting in a fixed *E*_*K*_. Dashed and dotted orange lines represent intermediate conditions where *K*_*o*_ increases faster than *K*_*a*_, producing depolarizing trajectories. The dashed and dotted red lines show the complementary case where *K*_*a*_ increases faster than *K*_*o*_, producing hyperpolarizing trajectories. The bottom and top axes indicate the corresponding values of *K*_*o*_ and *K*_*a*_, respectively.

According to our formulation, *V*_*a*_ finds its balance mostly between *V*_*s*_ (associated with the steady state of the syncytium) and *E*_*K*_. Therefore, we can safely assume that when *E*_*K*_ is more positive, *V*_*a*_ is pulled toward depolarization while maintaining a distance from *E*_*K*_ due to the isopotentiality effect from the syncytium [24, 47]. This distance will be reduced after MFA treatment due to the decrease of gap junction connections. Nevertheless, in both conditions, *V*_*a*_ will be depolarized. The least depolarized (dotted orange lines) and the most depolarized (dashed orange lines) curves have the same range of *K*_ext_ (*K*_*o*_: 5–10 mM), with a larger range over *K*_*a*_ for the dotted line (*K*_*a*_: 140–270 mM *<* 140–230 mM). Therefore, over the same rate of *K*_ext_ increments, a faster rising *K*_*a*_ (dotted line) leads to a slightly less depolarized *E*_*K*_.

Considering the reduction of gap junction permeability in the MFA condition, in principle, we should have a higher rate of intracellular K^+^ accumulation at steady state compared with control, which corresponds to the dotted and dashed orange lines observed here. Thus, while *V*_*a*_ maintains a certain distance from *E*_*K*_, it can still be pulled to a higher depolarized state, corresponding to what we observed in Figure 5, for MFA and control conditions.

It is important to note that we do not have access to the absolute values of membrane potentials in the experiments. However, under control conditions, the initial potential is in a hyperpolarized range, whereas after barium is applied to the bath, the experiment starts from a depolarized value. Therefore, when comparing depolarization magnitude in the presence of barium, it is more accurate to consider the percentage of depolarization relative to the increments of K^+^.

Moreover, as mentioned before, under the effect of barium, membrane potential does not reach steady state during the 20 seconds of the experimental protocol, which means that the dynamics are much slower compared to the other cases. Given enough time, at steady state, it is quite probable (as reported in previous works [4, 45]) that it can reach higher depolarization states than other conditions, which is aligned with the intuition that in the case of negligible K^+^ flux, the depolarization could be almost exclusively due to the increase of *K*_ext_. Thus, we can extrapolate that the green curve in Figure 5 could be a possible prediction for the barium condition at steady state.

#### 3.2.2 Kinetic properties and buffering efficiency

To analyze the membrane’s response to increased extracellular K^+^, we examined two key parameters: the slope of the rise and the time constant *τ*. Our model accurately reproduced the rising slope under all conditions and qualitatively matched the rise time constant (Figure 6).

**Figure 6:**
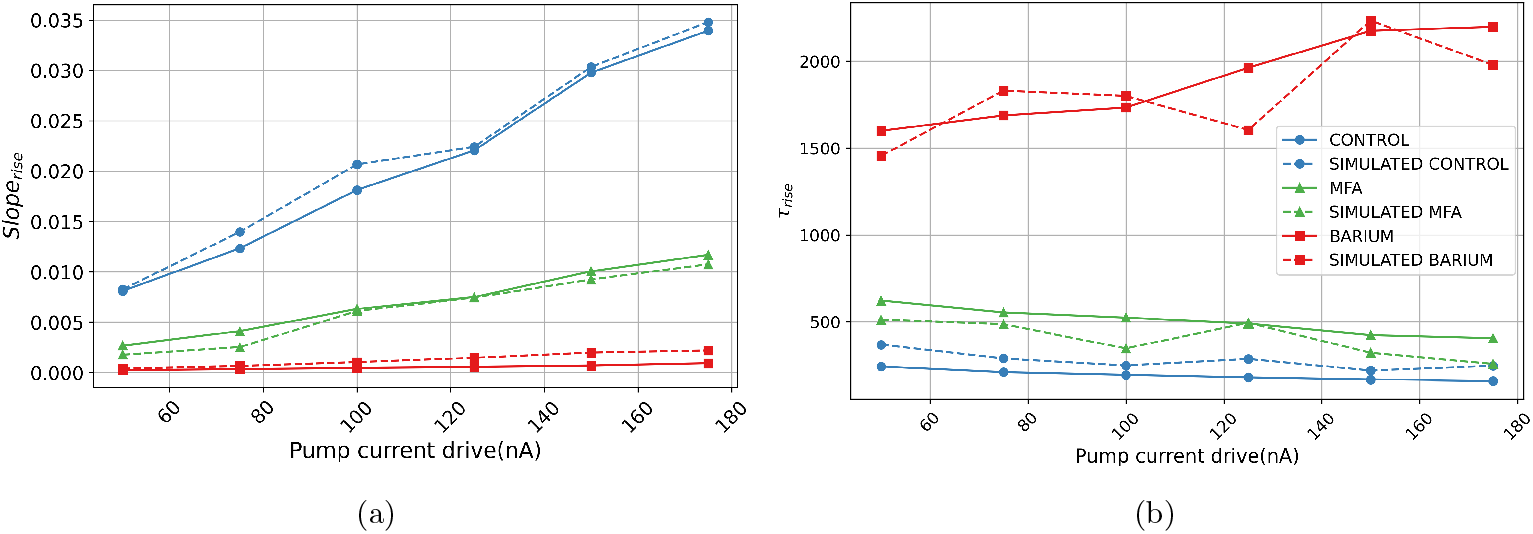
Dynamics of the Rise (a) Slope of Rise: This panel depicts the rate of change of membrane potential relative to iontronic pump current increments, showcasing a comparison between experimental values (solid lines) and simulated values (dashed lines). (b) Time Constant of Rise (*τ*_rise_): This plot illustrates the time constant of the membrane potential’s rise in response to iontronic pump current increments. It compares experimental data (solid lines) and simulations (dashed lines). Legends indicate the color codes corresponding to each experimental condition.

In the control condition, we observed the highest slope of rise, both in value and in the rate of increase over iontronic pump current increments. Under MFA treatment, the rate of change decreased, indicating slower dynamics during depolarization. With barium treatment, these metrics decreased dramatically, resulting in significantly slower dynamics (Figure 6a).

The time constant results further corroborated these observations (Figure 6b). In control conditions, we observed the lowest time constant (approximately 300 ms), which nearly doubled under MFA and increased up to 3 seconds under barium treatment. These observations indicate that faster dynamics correlate with higher buffering efficiency.

When examining the decay phase after stopping the iontronic pump (Figure 7), we observed that the control condition exhibited a rapid decline, the MFA condition a slower decline, and the barium-treated condition a very slow return to baseline. In all three cases, the rate of decline increased with successive iontronic pump increments, a trend that was qualitatively well reproduced by the simulated traces.

**Figure 7:**
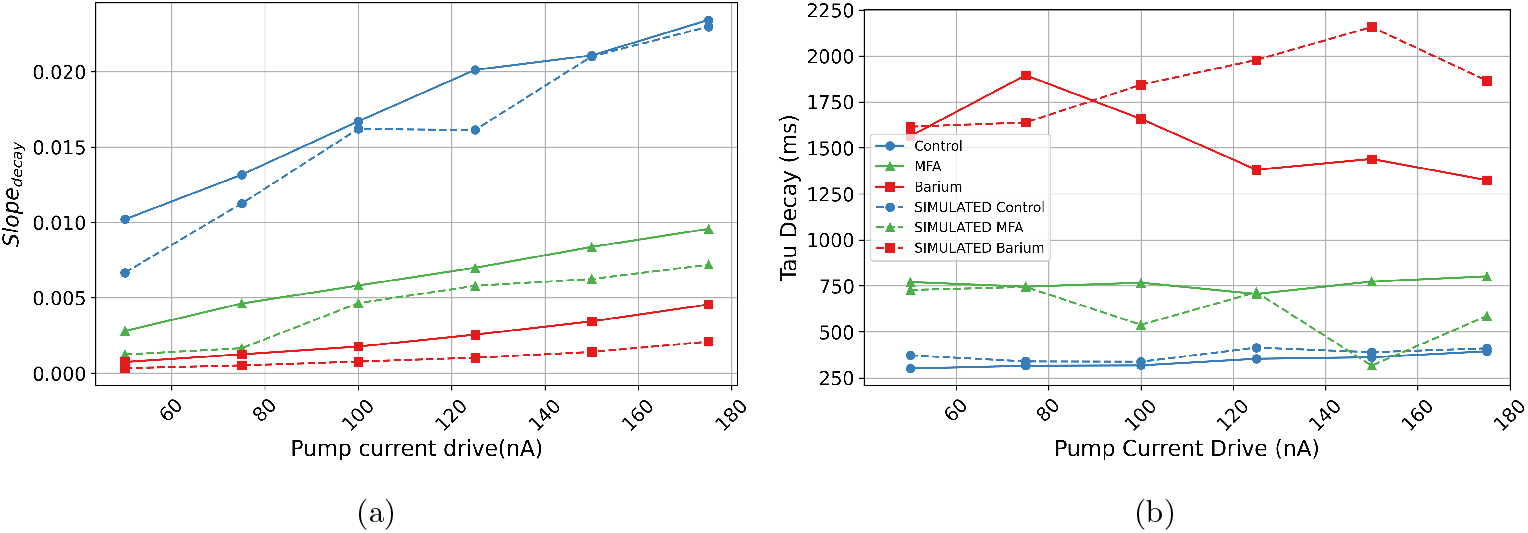
Dynamics of decay: (a) Slope of Decay: This panel depicts the rate of change of membrane potential decay, relative to iontronic pump current increments, showcasing a comparison between experimental values (solid lines) and simulated values (dashed lines). (b) Time Constant of Decay (*τ*_decay_): This plot illustrates the time constant of the membrane potential’s decay in response to iontronic pump current increments. It compares experimental data (solid lines) and simulations (dashed lines). Legends indicate the colour codes corresponding to each experimental condition.

### 3.3 Parameter space exploration and degeneracy

#### 3.3.1 Simulated data accounting for different iontronic pump currents

Our parameter optimization process explored the parameter space extensively, resulting in numerous successful model fits. Figure 8a presents 300 simulated membrane potential traces from the best-fitting parameter sets for the barium condition, all of which closely match the experimental recording.

**Figure 8:**
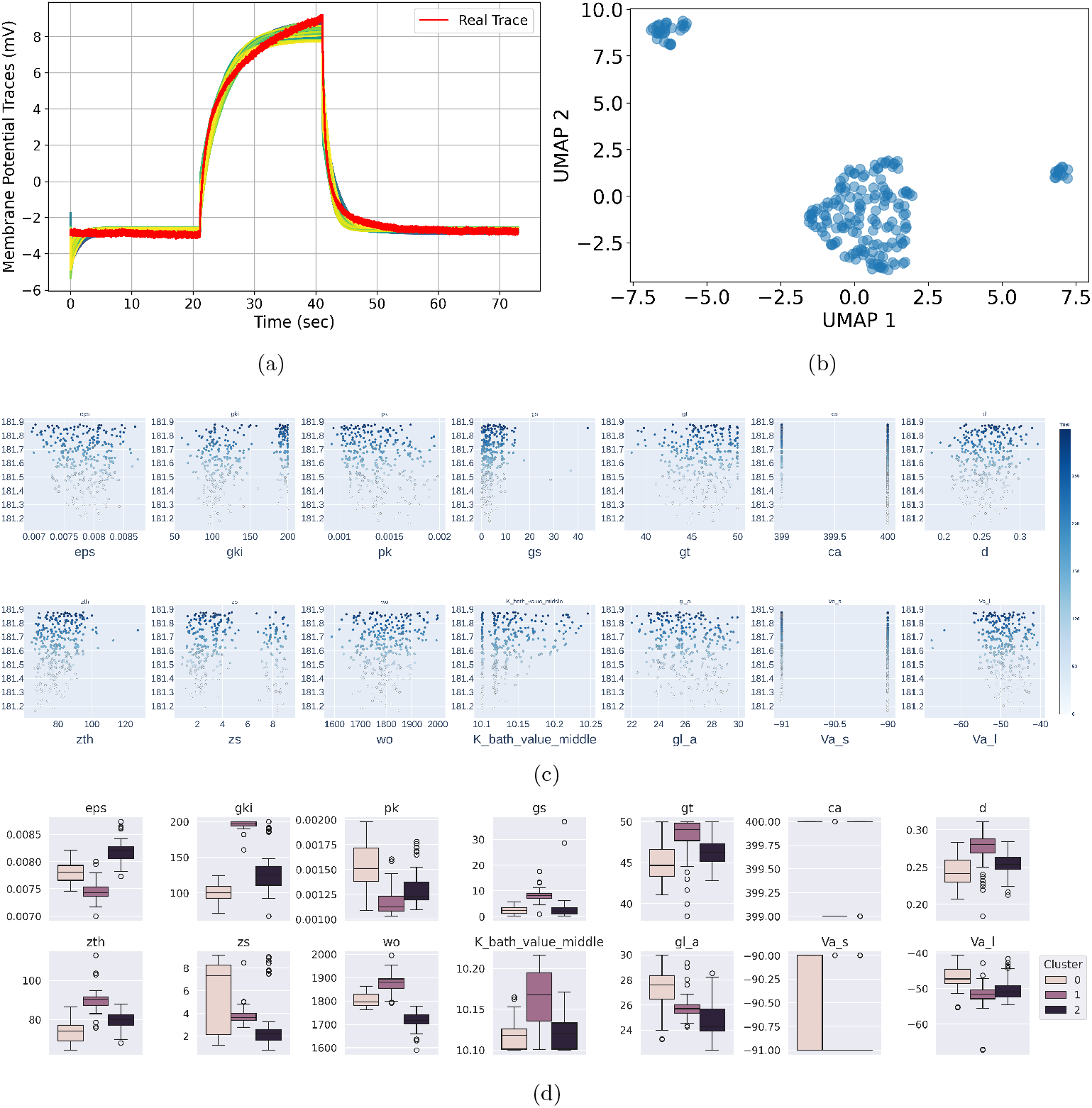
Degeneracy of model fits under the Barium condition. (a) Simulated membrane potential traces (yellow) from 300 final accepted trials of the optimization algorithm, overlaid on the experimental recording (red) for the 175 nA iontronic pump stimulation under Barium+MFA condition. These traces represent the subset of trials that yielded the lowest loss values and visually acceptable fits. (b) Slice plot for the 175 nA condition showing parameter values (x-axis) against their associated loss scores (y-axis, color-coded), indicating multimodal regions of good performance. (c) UMAP projection of 300 trials for the 150 nA condition, reduced from the full 14-dimensional parameter space to two dimensions and colored by unsupervised clustering. (d) Distribution of parameter values for the 150 nA condition shown across three clusters from (c), visualized as box plots. Each box represents the interquartile range per cluster and parameter; outliers are marked with dots. These panels collectively illustrate the model’s degeneracy—distinct parameter configurations producing similarly accurate fits—and support the inference of multiple biophysically plausible mechanisms underlying the Barium condition.

The distribution of optimized parameters revealed multimodal distributions when plotted against the objective score (Figure 8b), indicating degeneracy in our model—distinct parameter configurations can yield similar dynamic behaviors. To better understand the structure of the high-dimensional parameter space, we employed U-MAP dimension reduction (Figure 8c) and k-means clustering, which identified three clusters of parameter configurations.

The distribution of parameters across clusters (Figure 8d) showed that some parameters (*z*_th_, *ϵ*, and *g*_Ki_) had relatively well-defined boundaries, while others (*K*_bath_, *g*_La_, *d, g*_T_, and *g*_S_) exhibited substantial overlap between clusters. This pattern of distribution supports the presence of multimodality in the parameter space.

#### 3.3.2 Interaction Modality of Local and Spatial Buffering Parameters

Building on our distribution analysis, we applied spectral clustering to our best-fitting simulations (Figure 9). This analysis revealed three distinct clusters in the parameter space, particularly evident in the sigmoidal parameters that control spatial buffering. The clustering results suggest that the model’s behavior can be categorized into at least two primary modes, likely corresponding to different states of the sigmoidal function governing buffering processes. To mitigate potential sampling bias, we increased the number of trials and combined optimization data from all barium experiments. The observed multi-modality indicates model degeneracy, prompting further investigation.

**Figure 9:**
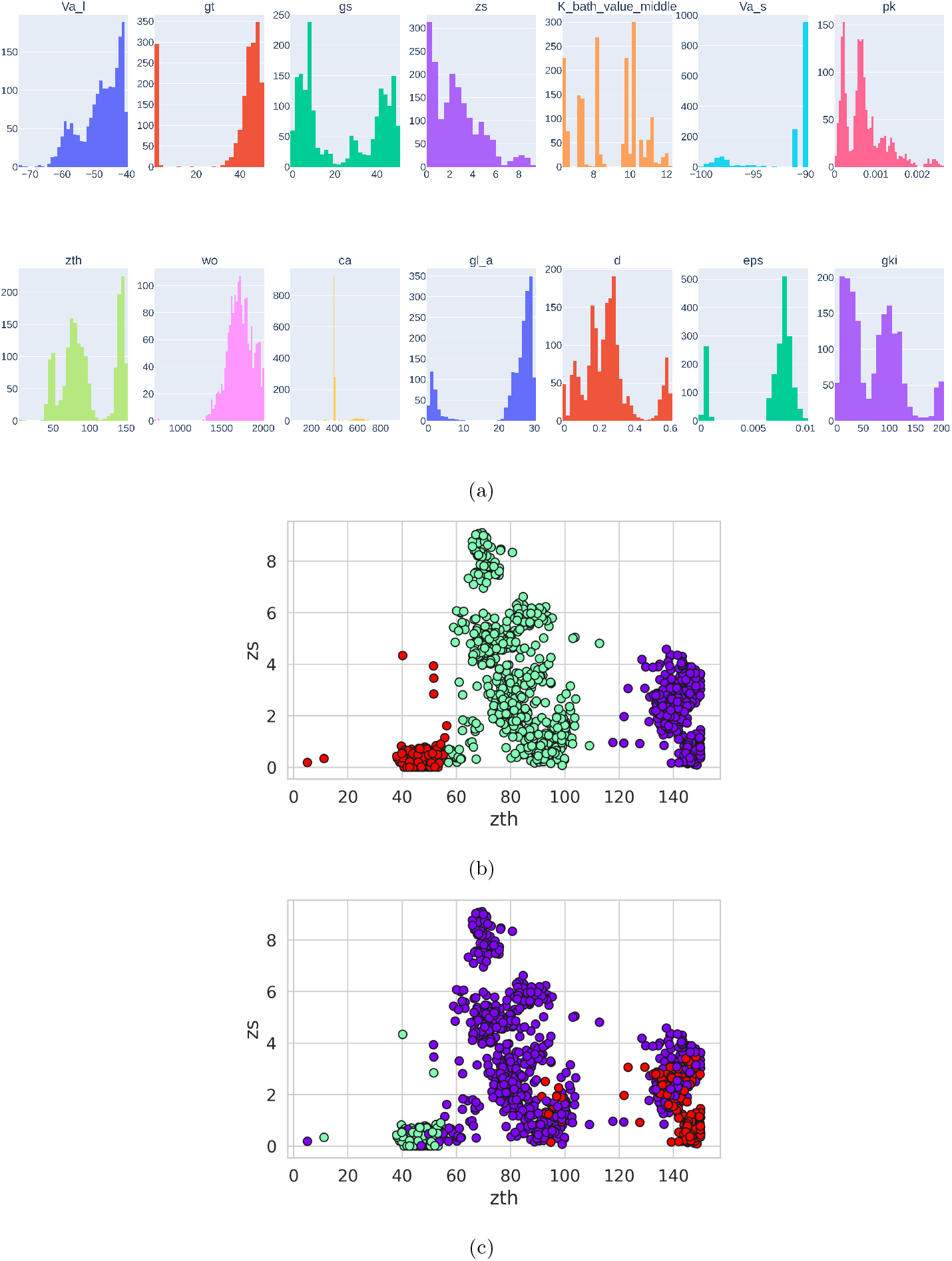
Modality due to the interplay of local and spatial buffering(a) Distribution histograms of parameter values from pooled top-performing trials across all barium experiment studies. The range of each parameter represents a subset of the predefined range of the optimization studies, shown across the pooled trials. The y-axis indicates the frequency of the loss function values or scores associated with each trial. (b): Spectral clustering visualization of the dataset with application of barium, with data points categorized into three distinct clusters based on their attributes for sigmoid parameters. Each color in the scatter plot represents one of the three clusters, determined using the nearest-neighbor method. This plot highlights the inherent grouping and diversity within the dataset across the selected features. (c): Spectral clustering of the dataset with application of barium with three clusters, performed using all available parameters but visualized specifically through the parameters *z*_*th*_ and *z*_*s*_. This scatter plot effectively demonstrates the clustering patterns based on the comprehensive dataset attributes while highlighting the distribution and interaction between *z*_*th*_ and *z*_*s*_. Each color in the plot represents one of the three distinct clusters, illustrating how these key parameters vary across the identified groups.

### 3.4 Degeneracy Supports Homeostatic Stability

To investigate whether our model demonstrates functional degeneracy over cell characteristics variability, we conducted simulations across a broad physiologically plausible range of K^+^ inputs (2.5–25 mM) with variable parameter configurations (Figure 10). These variations potentially reflect different astrocytic phenotypes across brain regions and physiological states.

**Figure 10:**
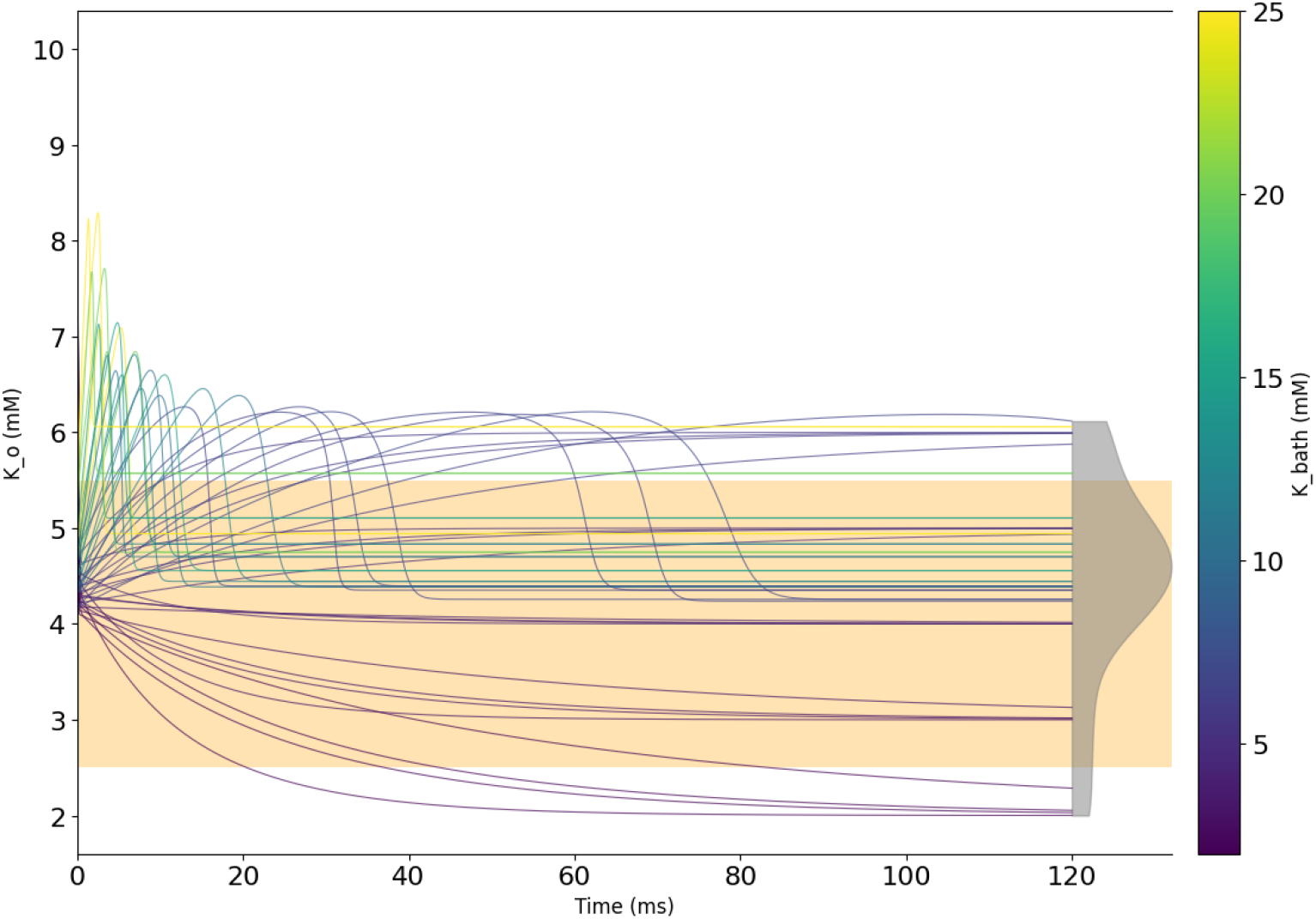
Simulated extracellular potassium concentration (*K*_*o*_) trajectories under different initial conditions and parameter configurations. Despite variations leading to undershoots and overshoots in *K*_*o*_, most traces converge at their steady-state to the same range (yellow shade), illustrating the model’s degeneracy in achieving potassium homeostasis.

Despite variations in individual parameters and initial conditions, the model consistently maintained stable extracellular K^+^ levels, displaying undershoots and overshoots before returning to steady-state levels within a physiological range (yellow shade in Figure 10). This stability highlights how diffusion, local buffering, and spatial buffering mechanisms compensate for each other to maintain system homeostasis.

The observation that most parameter configurations restore K^+^ homeostasis across an extensive range of inputs suggests that the homeostatic degeneracy of astrocytic K^+^ buffering is robustly represented by our model.

## 4 Discussion

### 4.1 Membrane Kinetics as a Window into Buffering Efficiency

To characterize the membrane’s response to elevated extracellular potassium ([*K*^+^]_ext_), we examined two kinetic parameters: the rate of change of membrane potential (slope) during the rise and decay phases, which reflects the speed of movement toward steady state, and the time constant *τ*, defined as the time required to reach one-third of the maximum amplitude.

Potassium influx into astrocytes is osmotically coupled with water entry, typically leading to an increase in cell volume under elevated [*K*^+^]_ext_ conditions [20]. However, our use of an iontronic K^+^ pump, as opposed to conventional KCl puffing methods, minimizes Cl^*−*^-mediated osmotic loading [16, 46]. This moderate K^+^ elevation is unlikely to cause significant volume changes. Furthermore, low-dose barium (100 *µ*M) has been shown not to affect astrocyte volume [43]. Therefore, we assume negligible changes in membrane surface area and capacitance across conditions (*C*_*m*_ ∝*A*), allowing us to omit volume effects from our analysis. Although any increase in capacitance would reinforce the deductions that follow, our framework remains grounded in a constant-capacitance model.

Under these assumptions, the slope of the membrane potential trace serves as a proxy for the total membrane current in somatic patch-clamp recordings, particularly during the transient to reach the steady state (as modeled by 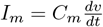). This approximation also allows estimation of total membrane conductance through the time constant relationship 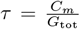. These assumptions are mostly valid in control and MFA conditions, where astrocytes are very sensitive to extracellular potassium dynamics and membrane potential shows strong correlation with [*K*^+^]_ext_ [8] (see also Fig. 3b–c).

Given this framework, the higher slope values observed during both depolarization and repolarization phases in control versus MFA (Figs. 6a, 7a) suggest higher total membrane current. This is corroborated by shorter time constants (Figs. 6b, 7b), indicative of higher conductance in control. This aligns with the known effect of MFA in reducing gap junction-mediated intracellular connectivity and thus the associated conductance.

This interpretation is further supported by Wallraff et al. [44], who demonstrated that astrocytes in hippocampal slices from CX30/CX43 double knockout (dKO) mice—lacking gap junction coupling— exhibited a ∼30% reduction in membrane current and increased membrane resistance. These observations mirror our findings under MFA treatment.

The relatively constant *τ* values across increasing K^+^ levels in both control and MFA suggest that membrane conductance remains approximately constant in these conditions. This can be explained by the limited recruitment of additional spatial connectivity (gap junctional connectivity/strength): in MFA due to gap junction blockade, and in control due to the moderate range of [*K*^+^]_ext_ elevation [2].

Given that conductance remains constant, the increasing slope across K^+^ increments in both control and MFA conditions (Fig. 6a) must primarily reflect an increase in the electrochemical gradient (*V*_*a*_−*E*_*K*_), driven by the progressive depolarization of *E*_*K*_ at higher extracellular K^+^. In control, this effect is further amplified, especially at higher K^+^ levels, by strong isopotentiality maintained through extensive gap junction coupling, which homogenizes the astrocytic membrane potential and holds *V*_*a*_ close to the network potential (*V*_*s*_ *< V*_*a*_ *< E*_*K*_) [24]. In MFA, despite impaired gap junction connectivity, the extended morphology of astrocytes still allows for a residual electrochemical gradient between *V*_*a*_ and *E*_*K*_, although this gradient is weaker compared to control. This morphological factor enables some degree of spatial potential coupling [22], which supports a moderate increase in slope at higher K^+^ levels, though less pronounced than in control.

Interestingly, this discrepancy in slope at higher K^+^ levels becomes less evident during the repolarization phase (Fig. 7a), when the reversal potential begins to hyperpolarize as extracellular K^+^ declines (Fig. 5). The resulting reduction in the electrochemical gradient (*E*_*K*_ → *V*_*a*_) may diminish the contribution of Kir-mediated spatial currents, as well as other passive K^+^ channels. Consequently, during decay, the slope difference between MFA and control exhibits a more linear trend across K^+^ increments.

This pattern is consistent with the observations of Larsen et al. [22], who reported that Kir-mediated spatial buffering contributes primarily during the rise phase of K^+^, whereas during decay, the dominant mechanism becomes the active Na^+^/K^+^-ATPase (NKA) pump.

Wallraff et al. [44] similarly reported that spatial buffering contributes most strongly to the fast initial phase of K^+^ clearance, particularly at high K^+^ levels, with this component being more pronounced in wild-type mice than in dKO mice. While it remains experimentally challenging to isolate the specific contribution of spatial buffering during the rise phase—due to overlapping mechanisms of activation— the more elevated [*K*^+^]_ext_ amplitude observed in dKO mice suggests that gap junction-mediated spatial buffering effectively contributes during the rise phase of K^+^, in agreement with [22].

Taken together, these observations support the hypothesis that the sharper depolarization rate in control compared to MFA (Fig. 6a) may serve as an indirect signature of enhanced Kir- and gap junction-mediated spatial buffering during the rise phase of K^+^ under high K^+^ conditions.

Further support comes from in vivo work by Chever et al. [8], who showed that astrocytes in glial-conditional Kir4.1 knockout (cKO) mice exhibit markedly slower membrane dynamics that fail to track changes in [*K*^+^]_ext_. They concluded that the dominant mechanism underlying astrocytic potential transients in these mice was the Na^+^/K^+^-ATPase (NKA). This phenotype closely resembles our observations under the combined MFA + barium condition, where both slope and *τ* metrics reveal impaired dynamics and the membrane potential fails to reach steady state during depolarization, unlike in control and MFA conditions alone.

Together, these studies link impaired K^+^ clearance with slowed astrocytic membrane kinetics, supporting the broader hypothesis that, under the conditions of this model, faster membrane dynamics are associated with more efficient K^+^ buffering.

Through systematic parameter searches and optimization, we found multiple distinct parameter sets that all produced similar astrocyte membrane dynamics. This underscores the inherent flexibility and robustness of astrocytic buffering. Rather than depending on a uniquely tuned set of parameters, astrocytes can achieve stable K^+^ homeostasis through a variety of configurations that reflect their natural variability. This capacity to generate comparable functional outcomes from diverse underlying conditions highlights the complexity and adaptability of astrocytic systems and supports the notion that stable network performance may arise from multiple distinct solutions. The properties of astrocytes can change in pathological conditions. Our results show that one cannot rely on measuring only one parameter, as changes in other parameters may generate a fully functional system following the degeneracy concept.

A key innovation of our approach lies in the introduction of a sigmoidal function that enables graded recruitment of spatial buffering in response to rising extracellular K^+^ levels. Unlike previous models that assume static buffering capacities or impose discrete thresholds, our formulation captures a more physiologically plausible and continuous transition between buffering regimes. This is inspired by experimental evidence that astrocytes modulate their buffering behavior depending on neuromodulatory state and extracellular K^+^ levels. Moreover, this flexibility may be shaped by cell volume changes, regional variations in syncytial coupling, or morphological differences, and may fluctuate over longer timescales such as circadian rhythms.

Our model, while capturing key aspects of astrocytic potassium buffering dynamics, remains highly simplified. By focusing on a minimal set of mechanisms, we have deliberately omitted other processes that are known to influence K^+^ homeostasis, including detailed Na^+^/K^+^-ATPase activity, glutamate-driven Na^+^ exchangers, and various metabotropic pumps and channels. Additionally, we treated the syncytium as an effectively steady-state compartment, which may not fully capture the spatial and temporal complexity of real astrocytic networks.

### 4.2 Conclusion

In this study, we provide three contributions to understanding astrocytic potassium buffering. First, we elucidate the role of spatial buffering in K^+^ regulation by linking experimental observations to the dynamics of the reversal potential. Second, we capture both fast and slow modes of K^+^ buffering, attributed to local and spatial mechanisms, without explicitly modeling sodium dynamics. Third, we demonstrate the presence of degeneracy in astrocyte membrane potential dynamics and homeostatic function, highlighting the system’s robustness and adaptability.

Our investigation reveals that astrocytes regulate K^+^ concentrations over multiple time scales through distinct buffering regimes. At lower levels of extracellular K^+^, the system relies predominantly on slow, local mechanisms governed by the Na^+^/K^+^-ATPase pump along with passive K^+^ channels and various exchangers. As K^+^ concentrations rise, a faster spatial buffering mode becomes prominent, driven by inward-rectifying potassium (Kir) channels working in concert with gap junctions to rapidly redistribute K^+^. Together, these local and spatial processes, along with the complex morphology of the astrocyte and the abundance of *K*^+^ leak channels, produce a wide spectrum of dynamics. The demonstrated degeneracy in both single-cell dynamics and homeostatic function highlights the robustness of this essential physiological process, which is critical to our understanding of resilience. Thus, it offers insight into how astrocytes may adapt to varying conditions while maintaining stable neuronal environments. By maintaining a minimal but flexible framework that can be systematically expanded, future models could account for more complex ionic exchanges, include more detailed neuronal components, and probe the conditions under which robust K^+^ homeostasis fails. Such models might help clarify how different pathways converge to produce similar dysfunctions, or how subtle changes in astrocytic buffering can influence global network excitability, offering valuable perspectives for understanding and eventually mitigating pathological states.

